# Chemoreceptor family in plant-associated bacteria responds preferentially to the plant signal molecule glycerol 3-phosphate

**DOI:** 10.1101/2024.12.10.627748

**Authors:** Félix Velando, Jiawei Xing, Roberta Genova, Jean Paul Cerna-Vargas, Raquel Vázquez- Santiago, Miguel A. Matilla, Igor B. Zhulin, Tino Krell

**Author notes:** Cold Spring Harbor Laboratory, Cold Spring Harbor, NY 11724, USA. These authors contributed equally to this work (Félix Velando, Jiawei Xing, Roberta Genova). Address correspondence to Igor Zhulin, or Tino Krell.

## Abstract

Plant pathogens and plant-associated bacteria contain about twice as many chemoreceptors as the bacterial average, indicating that chemotaxis is particularly important for bacteria-plant interactions. However, information on the corresponding chemoreceptors is limited. In this study, we identified the chemoreceptor PacP from the phytopathogen *Pectobacterium atrosepticum*, which exclusively recognized C3 phosphorylated compounds at its sCache ligand binding domain, mediating chemoattraction. Using a motif of PacP amino acid residues involved in ligand binding, we identified a chemoreceptor family, termed sCache_PC3, that was specific for C3 phosphorylated compounds. Isothermal titration calorimetry studies revealed that family members preferentially bound glycerol 3-phosphate, a key plant signaling molecule. Additionally, family members recognized glycerol 2-phosphate and glycolysis intermediates glyceraldehyde 3-phosphate, dihydroxyacetone phosphate and 3-phosphoglycerate. This study presents the first evidence of chemoreceptors that bind phosphorylated compounds. We show that the sCache_PC3 family has evolved from an ancestral sCache domain that respond primarily to Krebs cycle intermediates. Members of the sCache_PC3 family were mainly found in bacteria that interact with plants, including many important plant pathogens such as *Brenneria, Dickeya, Musicola, Pectobacterium,* and *Herbaspirillum*. Glycerol 3-phosphate is a signal molecule that is excreted by plants in response to stress and infection. Chemotaxis towards this molecule may thus be a means for bacteria to localize stressed plants and move to infection sites. This study lays the groundwork for investigating the functional importance of chemotaxis to phosphorylated C3 compounds in plant-bacteria interactions and virulence.

**Significance statement:** The bacterial lifestyle has shaped the evolution of signal transduction systems, and the number and type of chemoreceptors varies greatly between bacteria occupying various ecological niches. Our understanding of the relationship between lifestyle and chemoreceptor function is limited and the discovery of a chemoreceptor family in plant-associated bacteria that primarily responds to an important plant signal molecule is a significant advancement, allowing for further studies to determine its physiological relevance. The lack of knowledge about signals recognized by bacterial receptors is currently a major challenge in microbiology. This study illustrates the potential of combining experimental ligand screening with computational ligand prediction to identify signals recognized by uncharacterized receptors.

## Introduction

Chemotaxis is the directed active movement of bacteria in chemical gradients. The primary benefit of chemotaxis is the access to nutrients or the localization of sites that are favorable for survival (1). However, chemotaxis has also been observed to signals such as quorum sensing molecules, hormones and neurotransmitters that provide the bacterium with useful information about its microenvironment (2–5). Many bacteria establish interactions with other organisms, and chemotaxis to host compounds is frequently required for an efficient host colonization or virulence (6, 7).

Chemotaxis is typically initiated by signal binding at the ligand binding domain (LBD) of chemoreceptors. The chemotactic sensory capacity of a bacterium is reflected in the number of chemoreceptors that can differ greatly among bacteria, ranging from 1 to 90 (8). Bacterial lifestyle was found to determine the number of chemoreceptors (9, 10). Whereas bacteria that inhabit specific ecological niches possess few chemoreceptors, bacteria that live in variable environments or that maintain interactions with other living species have many more chemoreceptors. Chemoreceptors respond to many structurally different compounds including amino acids, organic and fatty acids, biogenic amines, polyamines, purine compounds, sugars, aromatic hydrocarbons, metal ions, inorganic anions, oxygen or polysaccharides (11). Although the link between chemoreceptor number and lifestyle has been established, we are still in the early stages of understanding the relationship between lifestyle and chemoreceptor function.

Chemotaxis is particularly important for the virulence of plant pathogens (6), where chemotaxis is frequently required for an efficient entry of bacteria into the host (12–14). Multiple lines of evidence indicate that compounds released by stomata and wounds attract bacteria to entry sites (6). The importance of chemotaxis in plant infection is further supported by the fact that phytopathogens and plant-associated bacteria have very broad chemosensory capabilities (15). On average phytopathogens possess 27 chemoreceptors, twice as many as bacteria that are not associated with plants (15). Many chemoreceptor families are almost exclusively present in phytopathogens, suggesting that they play specific roles in sensing plant compounds (15), however, revealing their specific functions awaits elucidation.

We use *Pectobacterium atrosepticum* as a model phytopathogen. It is among the 10 most relevant plant pathogens, causing black leg and soft rot diseases (16). The genome of *P. atrosepticum* strain SCRI1043 encodes 36 chemoreceptors that have a large variety of LBD types with different topologies (17). So far, three chemoreceptors that respond to quaternary amines, amino acids, and nitrate have been identified (18–20). The function of remaining 33 chemoreceptors remains unknown.

Cache domains comprise the largest superfamily of extracytosolic sensor domains in bacteria (21). They assume either a monomodular (single Cache domain, sCache) or bimodular configuration (double Cache domain, dCache). *P. atrosepticum* SCRI1043 has a single chemoreceptor with a sCache LBD (ECA_RS12390). Several sCache LBDs from chemoreceptors have been studied in phylogenetically diverse bacterial species, where they recognize different C1 to C4 carboxylic acids (22–27), and also urea and related compounds (28–30).

Over the last decade, the use of thermal-shift assay based ligand screening has become a very successful approach to identify the function of chemoreceptors (31–34). In addition, we have recently pioneered a computational approach that aids in the identification of receptor signals. Using LBD/ligand co-crystal structures and multiple alignments of homologous sequences, we derive sequence motifs that capture amino acid residues interacting with the ligand. Database searches for sequences that match these motifs has resulted in the identification of thousands of receptors that specifically bind amino acids (35), biogenic amines (36) and purines (37); results that were subsequently verified by Isothermal Titration Calorimetry (ITC) studies.

In the present study, we have combined both approaches. Thermal-shift assays and ITC studies of the LBD of ECA_RS12390 (termed PacP) revealed that it binds specifically phosphorylated C3 compounds including three glycolysis intermediates as well as glycerol 2-phosphate and glycerol 3-phosphate. Using a specific sequence motif present in the binding site, we define the corresponding domain family and confirm ligand-binding characteristics for selected members. Family members are almost exclusively present in plant-associated bacteria, including a number of important plant pathogens.

Glycerol 3-phosphate was preferentially recognized by the analyzed family members and induced the strongest chemotaxis in *P. atrosepticum*. Glycerol 3-phosphate is one of the most relevant plant signal molecules. For example, it regulates systemic immunity (38, 39) and responses to drought (40, 41). To the best of our knowledge, this is the first report on a bacterial receptor family that specifically binds phosphorylated compounds.

## Results

### Chemoreceptor ECA_RS12390 (PacP) binds exclusively phosphorylated C-3 compounds

To identify the ligands recognized by chemoreceptor ECA_RS12390, its individual LBD was overexpressed in *Escherichia coli* and purified by affinity chromatography. The purified LBD was subsequently subjected to thermal shift-based ligand screening assays using the compound arrays PM1, PM2A, PM3B and PM4A, which contain bacterial carbon, nitrogen, phosphorus, and sulphur sources (Fig. S1). This approach measures ligand-induced increases in the protein thermal stability, as quantified by the midpoint of the thermal unfolding transition (Tm). Significant increases were seen for glycerol 3-phosphate, carbamoylphosphate, and 3-phosphoglycerate among the 95 phosphorylated or sulfonated compounds of the PM4A compound array (Fig. 1A). No significant increases in Tm were observed for the remaining compound arrays, which include most of the carboxylic acids that were previously shown to bind to sCache_2 domains (22–27).

**Fig. 1).**
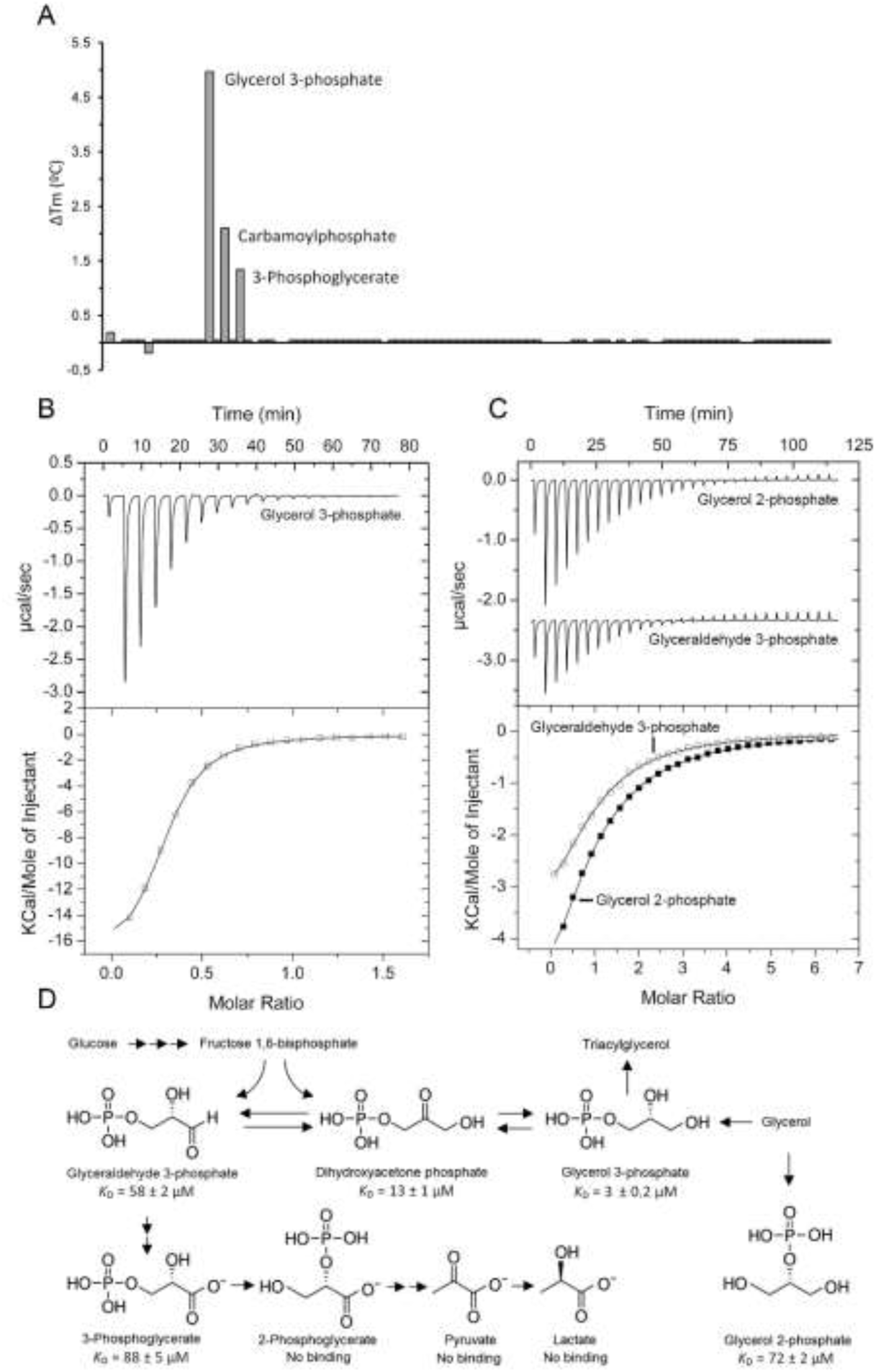
Binding of phosphorylated C3 compounds to the ligand binding domain of chemoreceptor ECA_RS12390 (PacP-LBD) **A)** Thermal shift assays with compounds of the Biolog array PM4A (phosphorus and sulfur sources). Shown are Tm changes with respect to the ligand-free protein. **B,C)** Microcalorimetric titration of 80 μM ECA_RS12390-LBD with 8 µl aliquots of 1 mM glycerol 3-phosphate, 2 mM glycerol 2-phosphate and 2.5 mM glyceraldehyde 3-phosphate. Upper panels: Titration raw data. Lower panels: Concentration-normalized and dilution heat-corrected integrated titration data. The lines are the best fits with the “One binding site model” of the MicroCal version of ORIGIN. **D)** Summary of ligands recognized by PacP-LBD and their metabolic relationships.

We subsequently conducted ITC binding studies to derive the dissociation constants (*K*_D_). The titration of ECA_RS12390-LBD with glycerol 3-phosphate resulted in large exothermic heat changes (Fig. 1B), leading to a calculated *K*_D_ of 3 ± 0.2 µM (Table 1). The binding of 3-phosphoglycerate occurred with significantly lower affinity (*K*_D_ = 88 ± 5 µM), whereas the titration with carbamoylphosphate did not result in measurable heat release. Because of the restriction of ligand dilution heats, microcalorimetry only permits monitoring of high-affinity binding events, indicating that carbamoylphosphate binds with an affinity that is not detectable by ITC.

**Table 1).** Dissociation constants derived from microcalorimetric binding studies of phosphorylated compounds to the LBDs of different chemoreceptors.

| Rec. No. | Accession code | Species | $K_D$ (μM) | | | | |
| --- | --- | --- | --- | --- | --- | --- | --- |
|  |  |  | Glyceraldehyde 3-P | Dihydroxyacetone P | 3-P glycerate | Glycerol 3-P | Glycerol 2-P |
| R1 | WP_011094075.1<br>(PacP, ECA_RS12390) | *P. atrosepticum* | $58 \pm 2$ | $13 \pm 1$ | $88 \pm 5$ | $3 \pm 0.2$ | $72 \pm 2$ |
| R2 | WP_136157342.1 | *Brenneria roseae* | $253 \pm 11$ | $99 \pm 10$ | $261 \pm 17$ | $11 \pm 0.2$ | $90 \pm 3$ |
| R3 | WP_006464688.1 | *Herbaspirillum frisingense* | Nb | $109 \pm 14$ | Nb | $33 \pm 1$ | $36 \pm 2$ |
| R4 | WP_158281851.1 | *Rivicola pingtungensis* | $100 \pm 2$ | $230 \pm 10$ | $926 \pm 8$ | $7 \pm 0.2$ | $60 \pm 2$ |
| R5 | WP_028444678.1 | *Chitinimonas koreensis* | Nb | Nb | Nb | Nb | Nb |
| R6 | WP_174062248.1 | *Agrobacterium larrymoorei* | Nb | Nb | Nb | Nb | Nb |
| R7 | WP_019915375.1 | *Methyloversatilis discipulorum* | $126 \pm 9$ | Nb | Nb | Nb | $223 \pm 45$ |
| R8 | WP_028536266.1 | *Paludibacterium yongneupense* | No protein overexpression |  |  |  |  |
| R9 | WP_189529961.1 | *Paludibacterium paludis* | Nb | Nb | Nb | Nb | Nb |
| R10 | WP_028866120.1 | *Psychromonas aquimarina* | Nb | Nb | Nb | Nb | Nb |

Nb: no binding

We next explored the binding of structurally related phosphorylated compounds that were not present in the compound arrays. No binding was observed for 2-phosphoglycerate, whereas glycerol 2-phosphate and glyceraldehyde 3-phosphate bound with *K*_D_ values of 72 ± 2 and 58 ± 2 µM, respectively (Fig. 1C, Table 1). In addition, dihydroxyacetone phosphate was recognized with a *K*_D_ of 13 ± 1 µM. Pyruvate and lactate, ligands recognized by other sCache LBDs (23–25, 42), did not bind, indicating that a phospho-moiety is required for binding. In summary, ECA_RS12390 binds specifically phosphorylated C3 compounds (Fig. 1D) and represents the first receptor that binds exclusively phosphorylated ligands.

Glyceraldehyde 3-phosphate, dihydroxyacetone phosphate, and 3-phosphoglycerate are glycolysis intermediates (Fig. 1D). Glycerol 3-phosphate is an intermediate that connects glycolysis, glycerol metabolism, and triacylglycerol synthesis (43, 44). Furthermore, glyceraldehyde 3-phosphate and 3-phosphoglycerate are intermediates of the Calvin–Benson–Bassham cycle that permits CO_2_ fixation in plants (45). We have renamed ECA_RS12390 PacP (*<u>P</u>ectobacterium <u>a</u>trosepticum* <u>c</u>hemoreceptor for <u>p</u>hosphorylated compounds).

### PacP mediates chemoattraction to phosphorylated compounds that bind to its LBD

We subsequently conducted quantitative capillary chemotaxis assays to determine whether PacP ligands induce chemotaxis. Modest chemoattraction was observed for glyceraldehyde 3-phosphate, dihydroxyacetone phosphate, and glycerol 2-phosphate. In contrast, chemotaxis toward glycerol 3-phosphate and 3-phosphoglycerate was significantly greater (Fig. 2).

**Fig. 2).**
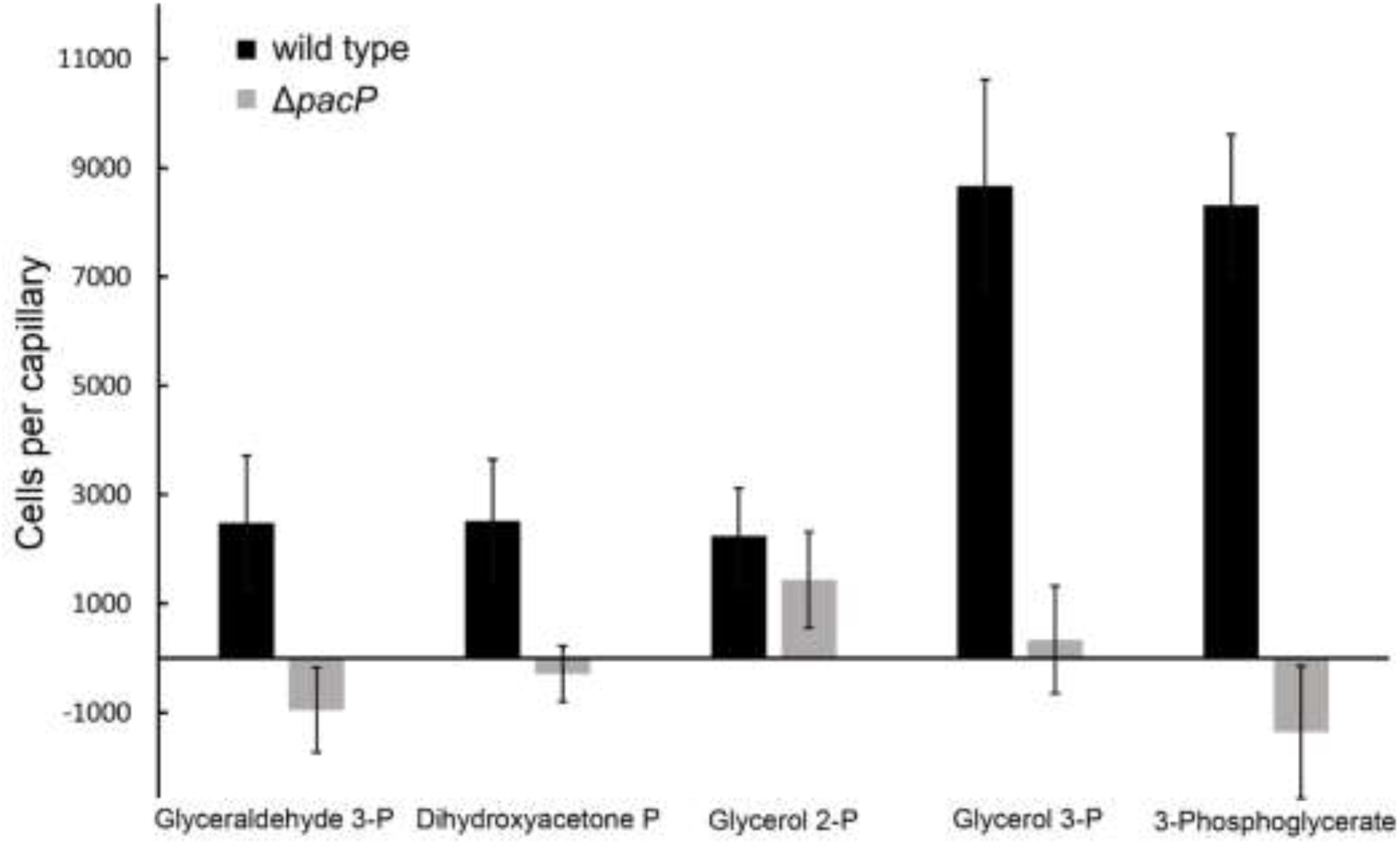
Capillary chemotaxis assays of *P. atrosepticum* SCRI1043 and a mutant deficient in *pacP* to 1 mM solutions of the ligands identified. Data have been corrected with the number of cells that swam into buffer-containing capillaries. Data are means and standard deviations from three biological replicates conducted in triplicate.

To assess the contribution of PacP in this response, we constructed a *pacP* mutant. Only glycerol 2-phosphate served as an attractant for the mutant, indicating that PacP is the sole chemoreceptor for glyceraldehyde 3-phosphate, dihydroxyacetone phosphate, glycerol 3-phosphate and 3-phosphoglycerate. An additional chemoreceptor(s) must exist for glycerol 2-phosphate.

### Three PacP ligands are of metabolic value

Chemoeffectors can be classified into three groups: compounds that are of metabolic value, compounds that act as environmental signaling cues, and compounds with a double metabolic/signal function (1, 46). To assess the metabolic value of the PacP ligands, we conducted growth experiments in minimal media containing the ligands as sole carbon or phosphorus source (Fig. 3).

**Fig. 3).**
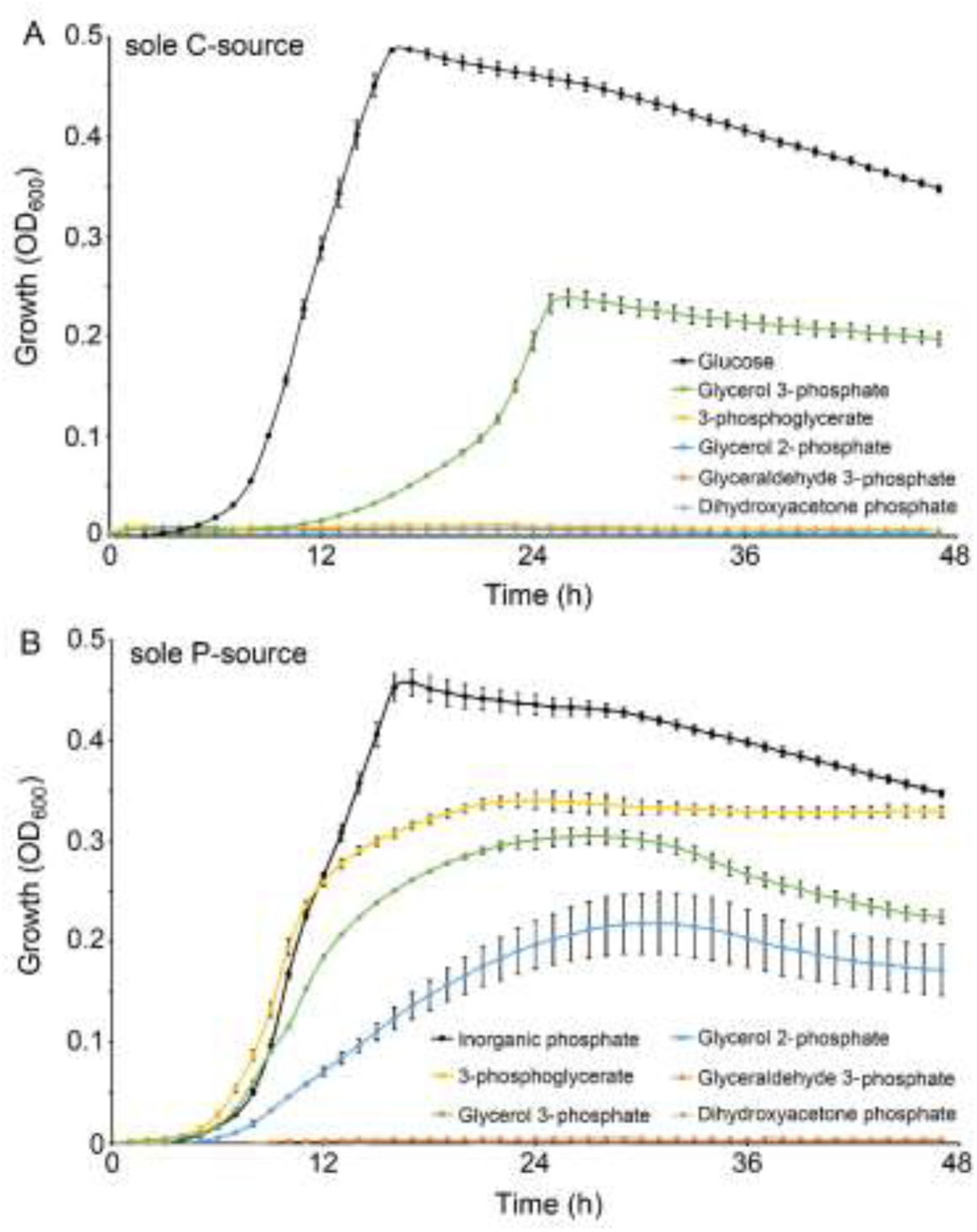
Growth of *Pectobacterium atrosepticum* SCRI1043 using the PacP ligands as sole carbon (A) or phosphorus (B) source. As reference, growth was monitored on glucose and inorganic phosphate as sole C- or P source. Data are means and standard deviations of three replicates.

Glycerol 3-phosphate was the only ligand that supported growth as sole carbon and phosphorus source, whereas 3-phosphoglycerate and glycerol 2-phosphate permitted growth as a phosphorus source. The glycolysis intermediates glyceraldehyde 3-phosphate and dihydroxyacetone phosphate were devoid of any apparent metabolic value.

### Definition of the sCache_PC3 family for phosphorylated compounds

Subsequent experiments were conducted to define the LBD family that binds phosphorylated C3 compounds. LBDs are rapidly evolving domains, and their ligand specificity is thus only poorly reflected in overall sequence identity (47). We have recently established a procedure to predict ligands recognized by sensor domains by taking into account the amino acid residues in the LBD binding site that interact with the bound ligand (35–37). Because extensive attempts to crystallize PacP-LBD failed, we constructed a model using AlphaFold2 (48) that was then used for computational ligand docking experiments.

These experiments indicated that Arg105 in PacP is likely the key residue for phosphate-sensing (Fig. 4A). A number of other amino acid residues, Y86, H100, Y121, K148, Y150, and Y167, also interacted with the bound ligand (Fig. 4A). To identify other chemoreceptors that may potentially bind phosphorylated compounds, we collected >1000 PacP homologs from the RefSeq protein database, from which 610 nonredundant sCache_2 domains were used to construct a phylogenetic tree (Fig. 4B). To probe which of these domains might recognize phosphorylated compounds similarly to PacP-LBD, we selected 10 domains from chemoreceptors from different clades (R1 to R10, Fig. 4B, C, Table 1) for further analyses. Their corresponding source strains were α-, β- and γ-proteobacteria that were isolated from plants, soil or freshwater (Table S1). Domains were selected for their pattern of conservation of residues within the predicted binding site (Fig. 4A) from the three distinct groups that are shown in different colors on Fig. 4B, C. Domains shown in red (including R1 to R6) have an invariant Arg105 (Fig. 4B, C). Domains shown in blue (including R7 and R8) have a Arg105Lys substitution (Fig. 4 B, C). Domains shown in grey, (including R9 and R10), have various other substitutions in the PacP position Arg105 (Fig. 4 B, C).

**Fig. 4).**
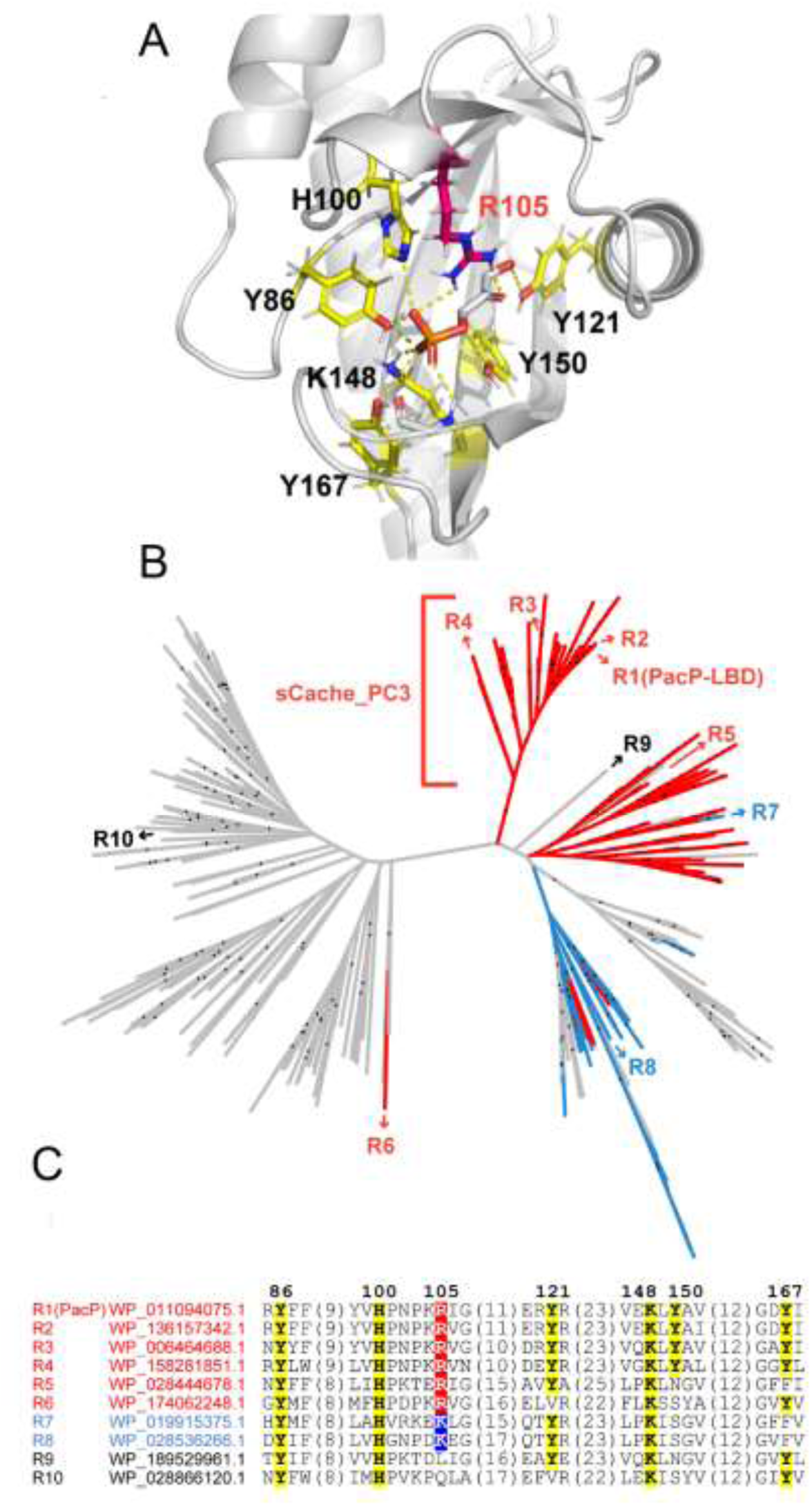
Definition of the sCache_PC3 family. **A)** PacP-LBD AlphaFold2 model containing docked dihydroxyacetone phosphate. Key residues are labelled. **B)** Maximum likelihood tree of sCache_2 domains from PacP homologs. Receptors R1 to R10, selected for experimental studies, are labeled. Dots indicate branches with bootstrap values not less than 70. **C)** Multiple sequence alignment of R1 to R10. The color of the protein name corresponds to that of panel B: red: conserved Arg105, blue: Lys instead of Arg105, grey: no Arg or Lys at position of Arg105. PacP residue numbering is shown.

To identify the ligands recognized by R2 to R10, expression plasmids harboring the codon-optimized LBD sequences were introduced into *E. coli*. Except for R8, all proteins could be overexpressed. Purified proteins were then subjected to microcalorimetric titrations. Receptors R2, R3, and R4 all bound phosphorylated compounds. Whereas, R2 and R4 bound all the five PacP ligands, receptor 3 bound only three of them: glycerol 3-phosphate, glycerol 2-phosphate, and dihydroxyacetone phosphate (Table 1). All four proteins bound glycerol 3-phosphate with the highest affinity, with dissociation constants ranging from 3 to 33 µM. These affinities are in the range typically observed for sensor protein-ligand interactions (49). Glycerol 2-phosphate was the second most tightly binding ligand for three proteins. The preferential recognition of glycerol 3-phosphate is illustrated by the binding studies of the 5 phosphorylated ligands to R4 (Fig. 5).

**Fig. 5).**
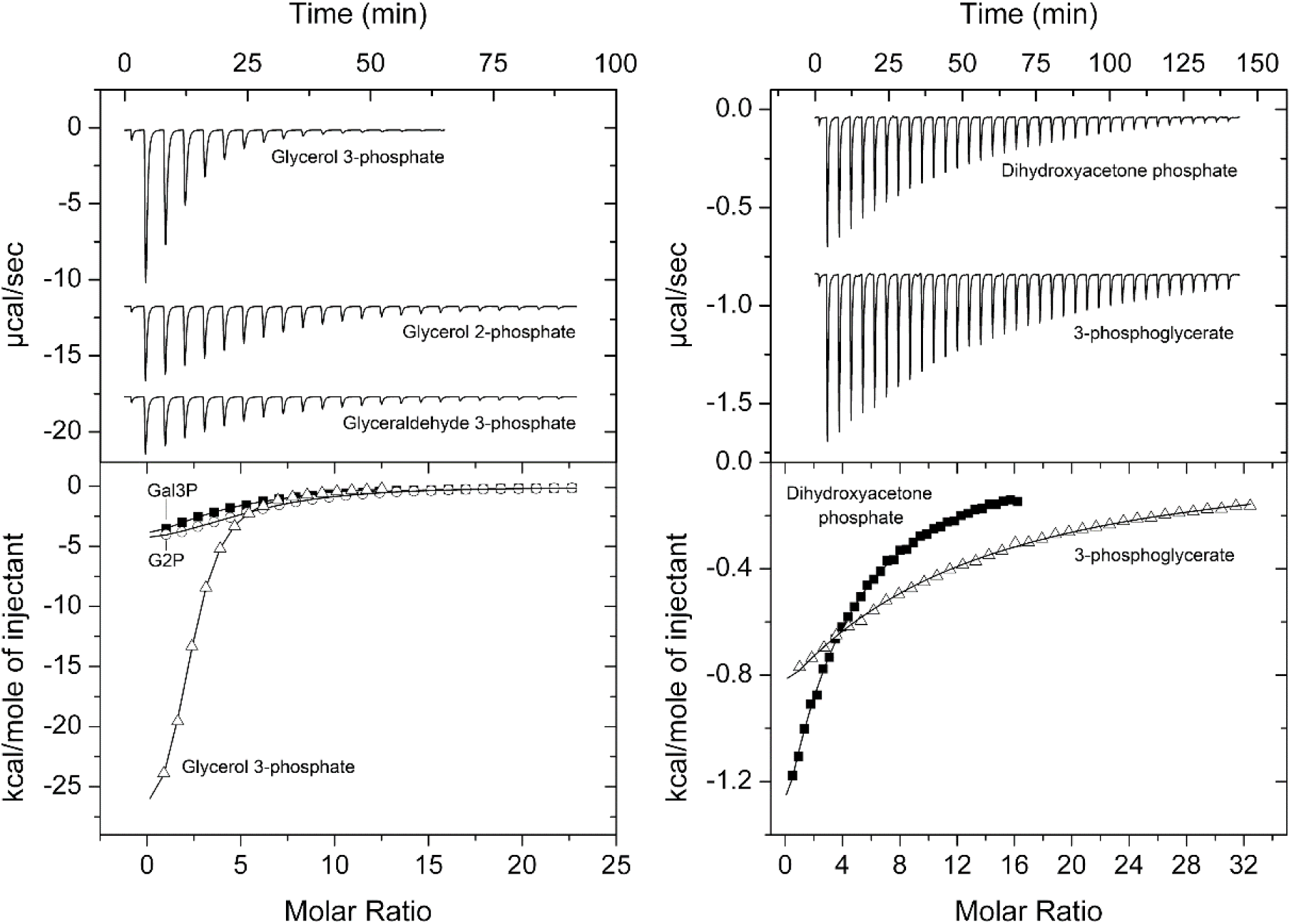
Microcalorimetric titration of the Ligand Binding Domain of R4 with different phosphorylated compounds. **A)** Titration of 33 µM protein with 8 µl aliquots of 3 mM glycerol 3-phosphate, 5 mM glycerol 2-phosphate (G2P) and 5 mM glyceraldehyde 3-phosphate (Gal3P). **B)** Titration of 33 µM protein with 8 µl aliquots of 2.5 mM dihydroxyacetone phosphate and 5 mM 3-phosphoglycerate. Upper panels: Titration raw data. Lower panels: Concentration-normalized and dilution heat-corrected integrated titration data. The lines are the best fits with the “One binding site model” of the MicroCal version of ORIGIN.

To determine whether receptors R2, R3, and R4 recognize other ligands apart from the phosphorylated compounds, we have conducted thermal shift assays with compound arrays PM1, PM2A, PM3B and PM4A (Fig. S1). Glycerol 3-phosphate was the only compound that caused significant increases in the Tm for proteins R3 and R4 (Fig. S2). Next to glycerol 3-phosphate, several small acids caused minor increases in the Tm of R2 (Fig. S2). However, ITC studies of these compounds to R2 showed binding exclusively for glycerol 3-phsophate. Taken together, data show that proteins R2, R3 and R4, like PacP, recognize exclusively phosphorylated C3 compounds.

Of the remaining proteins, only R7 bound phosphorylated compounds, namely glyceraldehyde 2-phosphate and glycerol 3-phosphate. However, the affinities were well below the affinities of the tightest binding ligands of receptors R1 to R4 (Table 1). These data indicate that the complete sequence motif present in R1 to R4 (Fig. 4C) is a prerequisite for high-affinity binding of phosphorylated compounds. The corresponding domain family has been termed sCache_PC3 (sCache domains for <u>p</u>hosphorylated <u>C3</u> compounds). The members of this family are provided in Data S1 and are exclusively found in chemoreceptors.

### The sCache_PC3 family has arisen from domains that bind different carboxylic acids

Because R5, R6, R9 and R10 failed to bind phosphorylated compounds, efforts were made to identify their ligands. To achieve this, purified proteins were analyzed by the thermal shift assay using the compound arrays PM1, PM2, PM3B and PM4. Compounds that caused significant Tm shifts were selected for microcalorimetric studies. All four receptors bound one or several of the Krebs cycle intermediates succinate, fumarate, malate or citrate (Fig. 6, Table 2). In addition, three domains bound other organic acids (Table 2), of which three, tricarballylate, methyl- and bromosuccinate, are structurally related to the Krebs cycle intermediates. The affinities of these compounds were in general lower than those of Krebs cycle intermediates (Table 2). Taken together, these findings suggest that domains others than those of the sCache_PC3 family (Fig. 4B) respond to Krebs cycle intermediates. The sCache_PC3 family likely evolved from these receptors found in *Burkholderiales* and *Enterobacterales*.

**Fig. 6).**
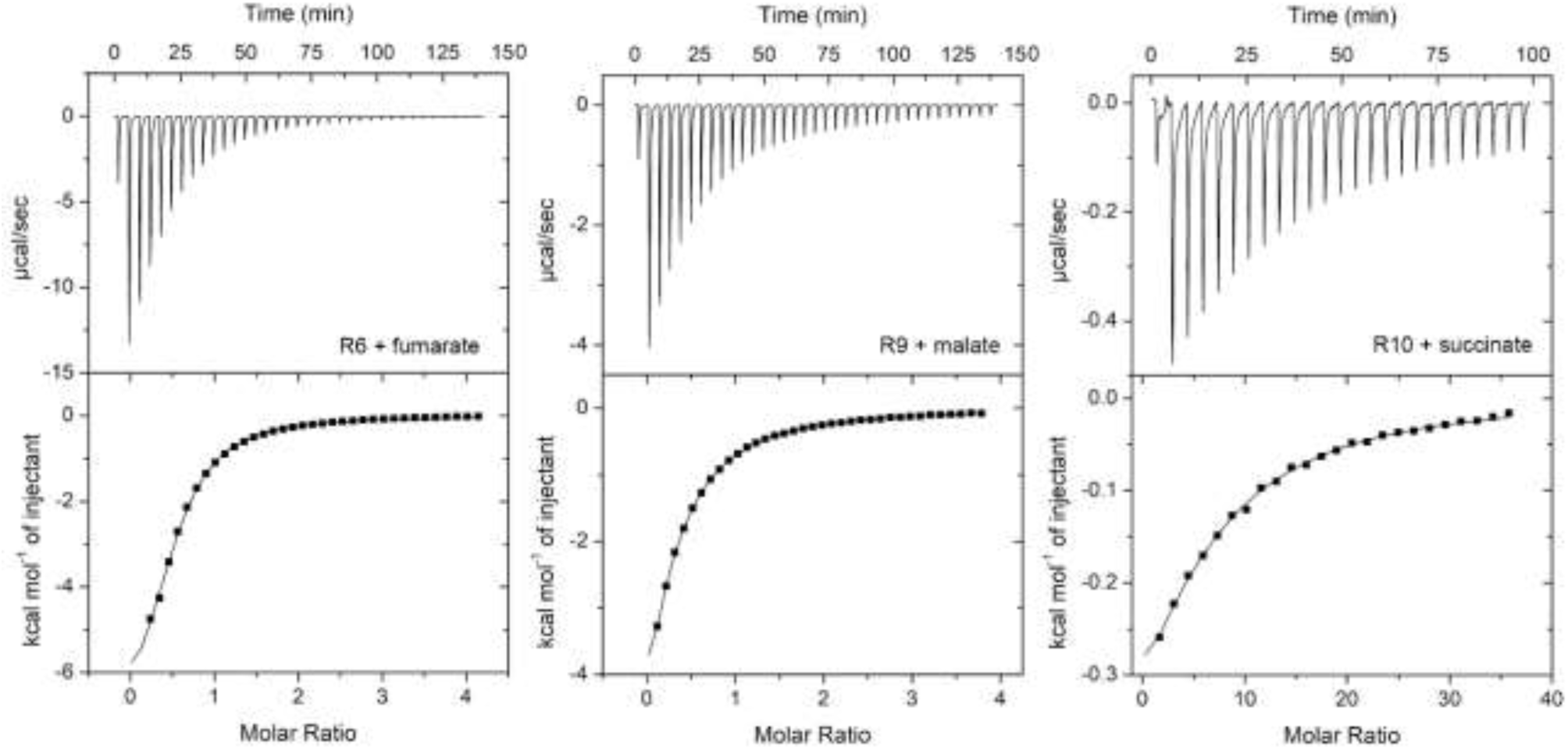
Microcalorimetric titrations of the ligand binding domain of different receptors that did not recognize phosphorylated compounds with Krebs cycle intermediates. Upper panels: Titration of 42 to 517 μM protein with 8 μl injections of 5 to 10 mM of different ligands. Lower panels: Concentration-normalized and dilution heat-corrected titration data. Dissociation constants are provided in Table 2.

**Table 2).** Dissociation constants derived from microcalorimetric binding studies of organic acids to the LBDs of different chemoreceptors.

| Rec.<br>No. | $K_D$ (μM) | | | | | | | | |
| --- | --- | --- | --- | --- | --- | --- | --- | --- | --- |
|  | Succinate | Fumarate | Malate | Citrate | Tricarallylate | Bromosuccinate | Methylsuccinate | Glycolate | L-lactate |
| R5 | $127 \pm 5$ | $68 \pm 6$ | $61 \pm 2$ | $10 \pm 2$ | $15 \pm 1$ | Nb | Nb | Nb | Nb |
| R6 | $90 \pm 4$ | $90 \pm 4$ | $63 \pm 1$ | Nb | Nb | Nb | Nb | Nb | Nb |
| R9 | Nb | Nb | $185 \pm 5$ | Nb | $138 \pm 33$ | $200 \pm 13$ | $254 \pm 53$ | Nb | Nb |
| R10 | $362 \pm 23$ | Nb | Nb | Nb | Nb | Nb | Nb | $322 \pm 114$ | $463 \pm 51$ |

Nb: No binding

### sCache_PC3 members are primarily present in plant-associated bacteria

Members of the sCache_PC3 family are chemoreceptors from γ-proteobacteria that belong to the orders Burkholderiales and Enterobacterales (Data S1). At the genus level the most abundant are *Pectobacterium*, followed by *Jantinobacterium, Acidovorax, Herbasprillum* and *Brenneria* (Fig. 7A). To gain insight into environments inhabited by these bacteria, we compiled the isolation sources of all family members (Data S1, Fig. 7B). Importantly, about two thirds of the strains were isolated from plants and another one quarter from freshwater and soil, which are two habitats in which plants are frequently found. In a number of cases strains of the same species have been isolated from plants and soil or freshwater (50, 51). Among the plant-associated strains, many belonged to species that have been associated with plant virulence such as *Brenneria*, *Dickeya*, *Musicola*, *Pectobacterium*, *Herbasprillum*, *Acidovorax*, and *Paracidovorax* (52, 53) (Data S1). Despite the very large number of strains that have been isolated from human and animals, only a few of them possess receptors with a sCache_PC3 domain (Fig. 7B), supporting the notion that sCache_PC3 containing chemoreceptors are predominantly present in plant-associated bacteria.

**Fig. 7).**
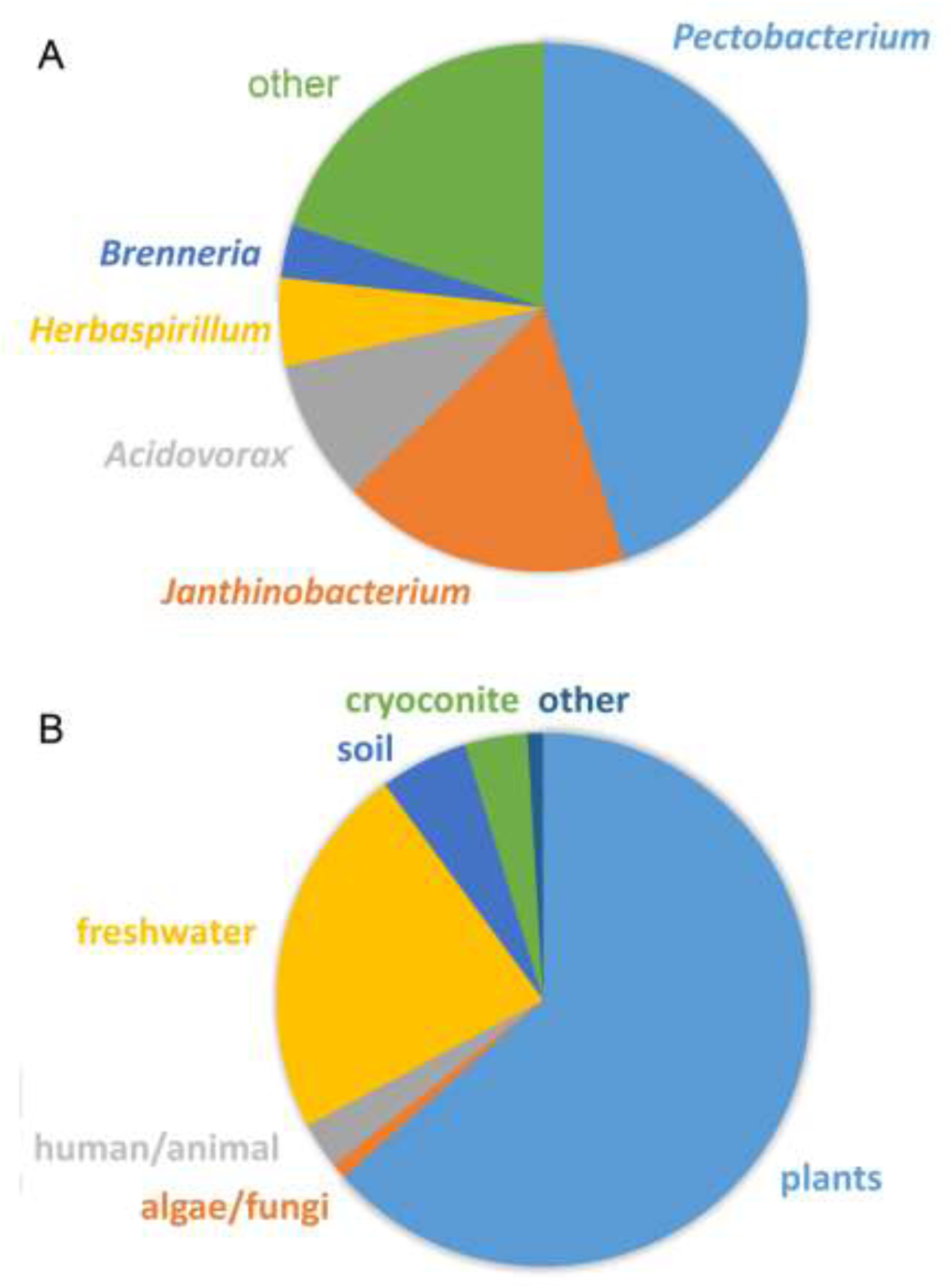
The sCache_PC3 family of sensor domains. **A)** Distribution of family members in different bacterial genera. Genera with less than 10 members are combined as “other”. **B)** Isolation sources of strains harboring chemoreceptors with sCache_PC3 domains. Detailed information is provided in Data S1.

## Discussion

The functions of flagellar motility and chemotaxis in bacteria are very diverse and depend on the bacterial lifestyles (46). Accessing nutrients and more generally locating niches that are optimal for growth appear the major functions of bacterial chemotaxis. Chemotaxis is also required for inter-domain communication permitting the establishment of beneficial or pathogenic interactions of bacteria with humans, animals and plants (6, 46). The diversity of functions of chemotaxis is reflected by the observation that bacteria of different lifestyles differ largely in the number and type of chemoreceptors (9, 10). Whereas bacteria that inhabit a specific ecological niche possess few chemoreceptors, bacteria that live in variable environments or that maintain interactions with other living species have many more chemoreceptors. However, since most chemoreceptors are of unknown function, our understanding of the link between chemoreceptor function and lifestyle is currently limited.

Plant pathogens and plant-associated bacteria stand out for their chemosensory capacities because they possess about twice as many chemoreceptors as the bacterial average (15). Many chemoreceptor families were found to be present primarily in plant-associated bacteria or phytopathogens (15). However, the ligands they recognize remain to a large degree unknown. The identification of a chemoreceptor family in plant-associated bacteria that responds to the plant signal glycerol 3-phosphate provides novel insight. In plants, glycerol 3-phosphate is a critical inducer of systemic acquired resistance (SAR) to pathogen attack; mutants defective in glycerol 3-phosphate synthesis are unable to induce SAR (38, 39, 54). Plant infection induces SAR by stimulating glycerol 3-phosphate biosynthesis via the upregulation of the glycerol 3-phosphate biosynthesis genes (55). Glycerol 3-phosphate is also abundantly present in the rhizosphere, since it was the only PacP ligand that could be detected in root exudates (56, 57).

What might be the physiological benefit of chemotaxis to glycerol 3-phosphate for plant pathogens? i) Our studies show that *P. atrosepticum* can use glycerol 3-phosphate as a sole carbon and phosphorus source for growth, and access to nutrients may be one of the benefits. ii) Glycerol 3-phosphate, which was found to accumulate at the site of infection (55), likely attracts bacteria to these locations, facilitating plant entry. iii) Glycerol 3-phosphate is a stress-related signal molecule, and its levels increase in response to phosphate starvation (56) and drought (40). Of the 114 compounds that were enriched in the drought-treated endosphere, glycerol 3-phosphate was increased about 22,000-fold. It was thus the compound with by far the most significant increase. Further experiments showed that this increase was due to changes in the host and not within the rhizosphere (40). Because it is a stress signal, chemotaxis to glycerol 3-phosphate may also be a means to locate and move to stressed plants that are an easier target for infection than non-stressed plants. Glycerol 3-phosphate is also an important bacterial signal molecule. For example, in *P. aeruginosa*, glycerol 3-phosphate accumulation affects twitching motility, pyocyanine and exopolysaccharide biosynthesis, antibiotic resistance, and tolerance to oxidative stress (58). Further studies will show to what degree glycerol 3-phosphate impacts the physiology of plant-associated bacteria.

The lack of information on signals that stimulate bacterial receptors is currently a major limitation in microbiology representing an important research need. Here we have combined two experimental strategies that were successfully used in the past to identify receptor signals, namely thermal shift based high-throughput ligand screening (31, 33, 34) and the use of binding pocket amino acid motifs derived from LBD-signal 3D structures (35–37). In contrast to the latter three studies, that were based on experimental LBD-signal co-structures, we identified a sequence motif in a ligand-docked AlpaFold2 model of PacP-LBD. This approach does thus not require experimental protein structures, suggesting that it might be universally applicable to identify receptor signals. The sCache_PC3 family is characterized by the Y(13)H(4)R(15)Y(26)K(1)Y(16) sequence motif (Fig. 4C) that can be used to identify further family members. Combining these two strategies is thus a very potent approach to identify novel sensor domain families providing insight into receptor function. This approach is not restricted to chemoreceptors and can be used to identify signals for LBDs of any other receptor family.

Chemotaxis has so far been observed for many compound families including amino-, organic, and fatty acids, sugars, polyamines, quaternary amines, purines, pyrimidines, aromatic hydrocarbons, oxygen, inorganic ions, and polysaccharides (11, 59). This is the first report of chemoreceptors that recognize specifically phosphorylated compounds, expanding our knowledge on the sensory capacity of chemoreceptors. Previous studies reported weak chemotaxis of *Bdellovibrio bacteriovorus* and *Bacillus subtilis* to 3-phosphoglycerate, glycerol 3-phosphate and glycerol 2-phosphate (60, 61). However, these bacteria do not possess sCache_PC3 containing chemoreceptors (Data S1), indicating that there are other types of chemoreceptors that mediate such responses. To the best of our knowledge, this is the first report of chemotaxis to the glycolysis intermediates glyceraldehyde 3-phosphate and dihydroxyacetone phosphate.

The identification of the sCache_PC3 family is a significant contribution to the field of plant-microbe interactions. This study forms the basis for further studies to explore to physiological relevance of bacterial chemotaxis to phosphorylated C3 compounds. It will also serve as an experimental guide to define other domain families.

## Materials and Methods

### Bacterial strains and growth conditions

Bacterial strains used in this study are listed in Table S2. *P. atrosepticum* SCRI1043 and its derivative strains were grown at 30 °C in Luria Broth (5 g/l yeast extract, 10 g/l bacto tryptone, 5 g/l NaCl) or minimal medium (0.41 mM MgSO_4_, 7.56 mM (NH_4_)_2_SO_4_, 40 mM K_2_HPO_4_, 15 mM KH_2_PO_4_) supplemented with 0.2 % (w/v) glucose as carbon source. *E. coli* strains were grown at 37 °C in LB. When appropriate, antibiotics were used at the following final concentrations (in μg/ml): kanamycin, 50; ampicillin, 100; streptomycin, 50.

### Plasmids and mutants

A mutant defective in *ECA_RS12390* was constructed by homologous recombination using a derivative plasmid of the suicide vector pKNG101. Briefly, a 0.5-kb fragment corresponding to the region encoding the LBD of ECA_RS12390 was amplified using primers specified in Table S2 and cloned into pGEM^®^-T. The resulting plasmid pGEM::ECA_RS12390 was digested with the XmaI and the 0.5-Kb insert was cloned into the same site at pKNG101 to generate pKNG101::ECA_RS12390. This plasmid was transferred into *P. atrosepticum* SRCI1043 by biparental conjugation using *E. coli* β2163. All plasmids were verified by PCR and sequencing. The pET28b(+) derivatives encoding the LBD of the different receptors analyzed in this study (Table 1) were purchased from GeneScript. The sequences of the proteins analyzed in this study are provided in Table S3.

### Protein overexpression and purification

Plasmids for the overexpression of the different proteins were transformed into *E. coli* BL21(DE3). The resulting strains were grown under continuous stirring (200 rpm) at 30 °C in 2-liter Erlenmeyer flasks containing 500 ml LB medium supplemented with 50 μg/ml kanamycin. At an OD_660_ of 0.5, protein expression was induced by the addition of 0.1 mM isopropyl β-D-1-thiogalactopyranoside. Growth was continued at 16 °C overnight prior to cell harvest by centrifugation at 10 000 x *g* for 20 min. Cell pellets were resuspended in buffer A (Table S4) and subsequently broken by French press treatment at 62.5 lb/in^2^. After centrifugation at 20 000 x *g* for 30 min, supernatants were loaded onto 5-ml HisTrap HP columns (Amersham Biosciences) equilibrated with buffer A and proteins eluted with a linear gradient of buffer B (Table S4). All proteins were purified at 4 °C. Purified proteins were dialyzed overnight into the corresponding analysis buffers (Table S4) for immediate analysis.

### Differential Scanning Fluorimetry-Based Thermal Shift Assays

Assays were carried out using a MyIQ2 Real-Time PCR instrument (BioRad, Hercules, CA, USA). Ligand solutions were prepared by dissolving the array compounds in 50 µl of MilliQ water, which, according to the manufacturer, corresponds to a concentration of 10–20 mM. Freshly purified proteins were dialyzed into the analysis buffer (Table S4). Compound arrays PM1, PM2A (carbon sources), PM3B (nitrogen sources) and PM4A (phosphorus and sulphur sources) from Biolog (https://www.biolog.com/) were used. The composition of these arrays are provided in Fig. S1. Experiments were conducted in 96-well plates and each assay mixture contained 20 µl of the dialyzed protein (at 80–50 µM), 2 µl of 5× SYPRO orange (Life Technologies, Eugene, Oregon, USA) and 2.5 µl of the resuspended array compound or the equivalent amount of buffer in the ligand-free control. Samples were heated from 23 to 85 °C using a scan rate of 1 °C/min. The protein unfolding curves were monitored by detecting changes in SYPRO Orange fluorescence. The Tm values were determined using the first derivative values of the raw fluorescence data.

### Isothermal titration calorimetry (ITC)

All experiments were conducted on a VP-microcalorimeter (Microcal, Amherst, MA) at 15 °C for PacP-LBD and at 20 °C for the remaining proteins. Proteins were dialyzed into the analysis buffers specified in Table S4, placed into the sample cell and were titrated with aliquots of ligand solutions (1 to 10 mM) freshly prepared in dialysis buffer. The mean enthalpies measured from the injection of the ligands into analysis buffer were subtracted from raw titration data prior to data analysis with the MicroCal version of ORIGIN. Data were fitted with the ‘One binding site model’ of ORIGIN.

### Quantitative capillarity chemotaxis assays

Overnight cultures of *P. atrosepticum* SCRI1043 were grown at 30 °C in minimal medium. At an OD_660_ of 0.4-0.45, the cultures were washed twice with chemotaxis buffer (50 mM K_2_HPO_4_/KH_2_PO_4_, 20 μM EDTA, 0.05% (v/v) glycerol, pH 7.0) and diluted to an OD_660_ of 0.1 in the same buffer. Subsequently, 230 μl of the resulting bacterial suspension were placed into the wells of 96-well plates. One-microliter capillary tubes (P1424, Microcaps; Drummond Scientific) were heat-sealed at one end and filled with either the chemotaxis buffer (negative control) or chemotaxis buffer containing PacP ligands at a concentration of 1 mM. The capillaries were immersed into the bacterial suspensions at its open end. After 30 min at room temperature, the capillaries were removed from the bacterial suspensions, rinsed with sterile water and the content expelled into 1 ml of 40 mM K_2_HPO_4_, 15 mM KH_2_PO_4_. Serial dilutions were plated onto minimal medium supplemented with 15 mM glucose as carbon source. The number of colony forming units was determined after 36 h incubation at 30 °C. In all cases, data were corrected with the number of cells that swam into buffer containing capillaries.

### Growth experiments

*P. atrosepticum* SCRI1043 was cultured overnight in Phosphate-free M9-based medium (PFM9) (62) supplemented with 20 mM glucose as carbon source and 500 µM Pi as phosphate source. Cultures were washed and then diluted to an OD_600_ of 0.02 in PFM9 supplemented with different PacP ligands as sole carbon or phosphorous source at a final concentration of 5 mM. Subsequently, 200 µl aliquots of these cultures were transferred into microwell plates, and growth at 30°C was followed on a Bioscreen microbiological growth analyzer (Oy Growth Curves Ab Ltd., Helsinki, Finland) for 48h.

### Bioinformatic analyses

Molecular docking was performed on PacP and phosphorylated compounds using DiffDock with 10 inference steps (63). Homologs of the PacP (WP_011094075.1) N-terminal fragment (residues 7-197) were collected from RefSeq using BLAST (E-value < 0.05) (64). Sequences were aligned using MAFFT (65) and reduced to 98% redundancy using CD-HIT (66). The sCache_2 domain regions from the aligned sequences (corresponding to residues 27-178 in PacP) were used for phylogenetic analyses (67). A maximum likelihood tree was constructed using MEGA with the JTT model and 100 bootstraps (68). The protein targets were selected based on their location on the phylogenetic tree and the presence of key residues for ligand-binding.

## Acknowledgements

This work was supported by the Ministerio de Ciencia e Innovación /Agencia Estatal de Investigación 10.13039/501100011033 (grants PID2020-112612GB-I00 and PID2023-146216NB-I00 to TK, grants PID2019-103972GA-I00 and PID2023-146281NB-I00 to MAM), the Consejería de Economía, Innovación, Ciencia y Empleo, Junta de Andalucía (grant P18-FR-1621 to TK), the CSIC (grant 2024AEP062 to TK) and the US National Institutes of Health (grant R35GM 131760 to IBZ). JPCV was supported by the grant Unión Europea-NextGeneration EU RD 289/2021 UPM-Recualifica Margarita Salas.

## Abbreviations

ITC: isothermal titration calorimetry;
LBD: ligand binding domain;
SAR: systemic acquired resistance

## Author contributions

Conceptualization: M.A.M., I.B.Z. and T.K.; bioinformatic analyses: J.X.; investigation and interpretation: F.V., J.X., R.G., J.P.C.V. and R.V.S.; writing: I.B.Z., T.K. and M.A.M.; supervision: I.B.Z., T.K. and M.A.M.

## Materials & Correspondence

Correspondence and requests for materials can be sent to Igor B. Zhulin or Tino Krell.

## Supplementary Material

**Fig. S1).**
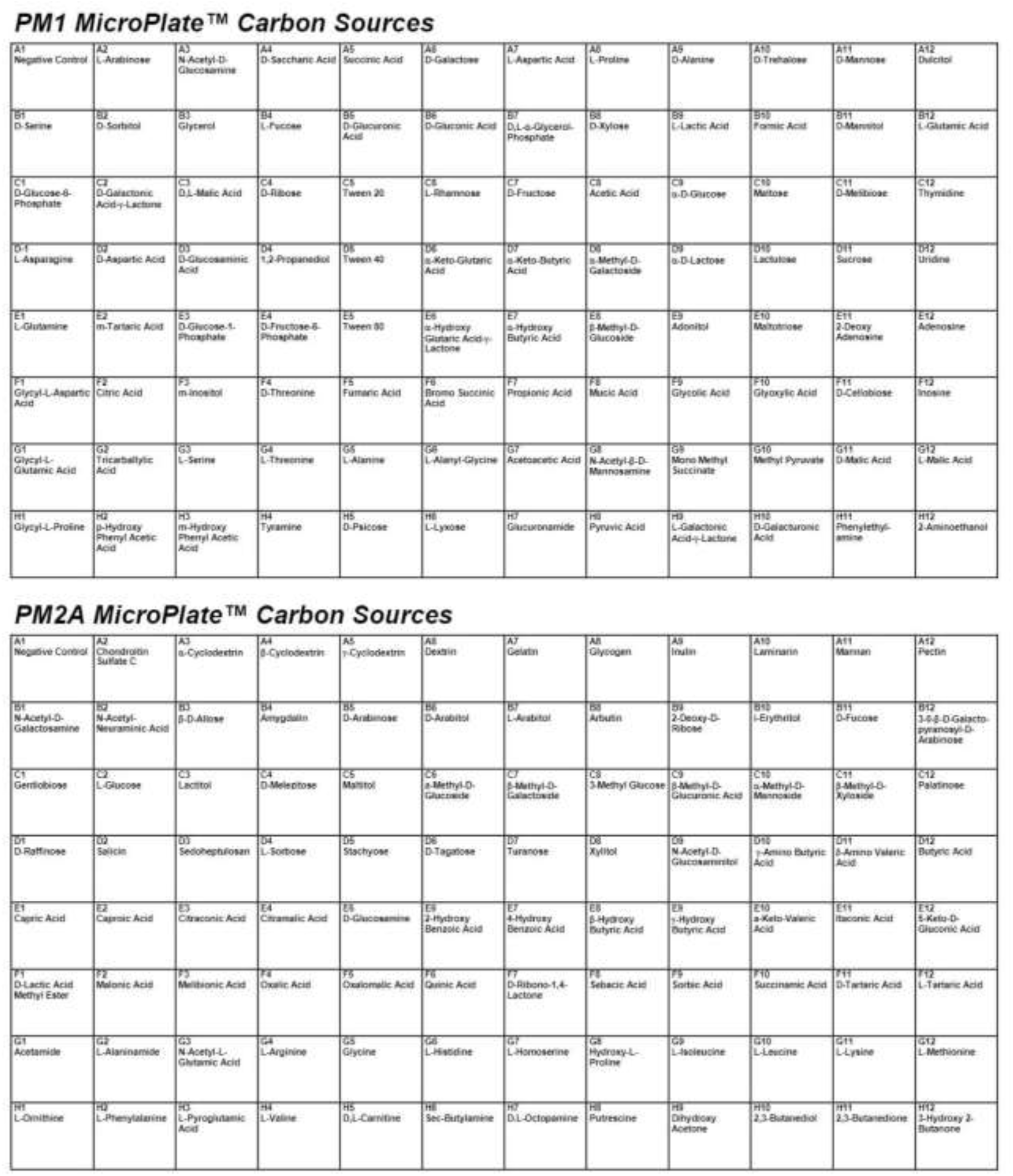
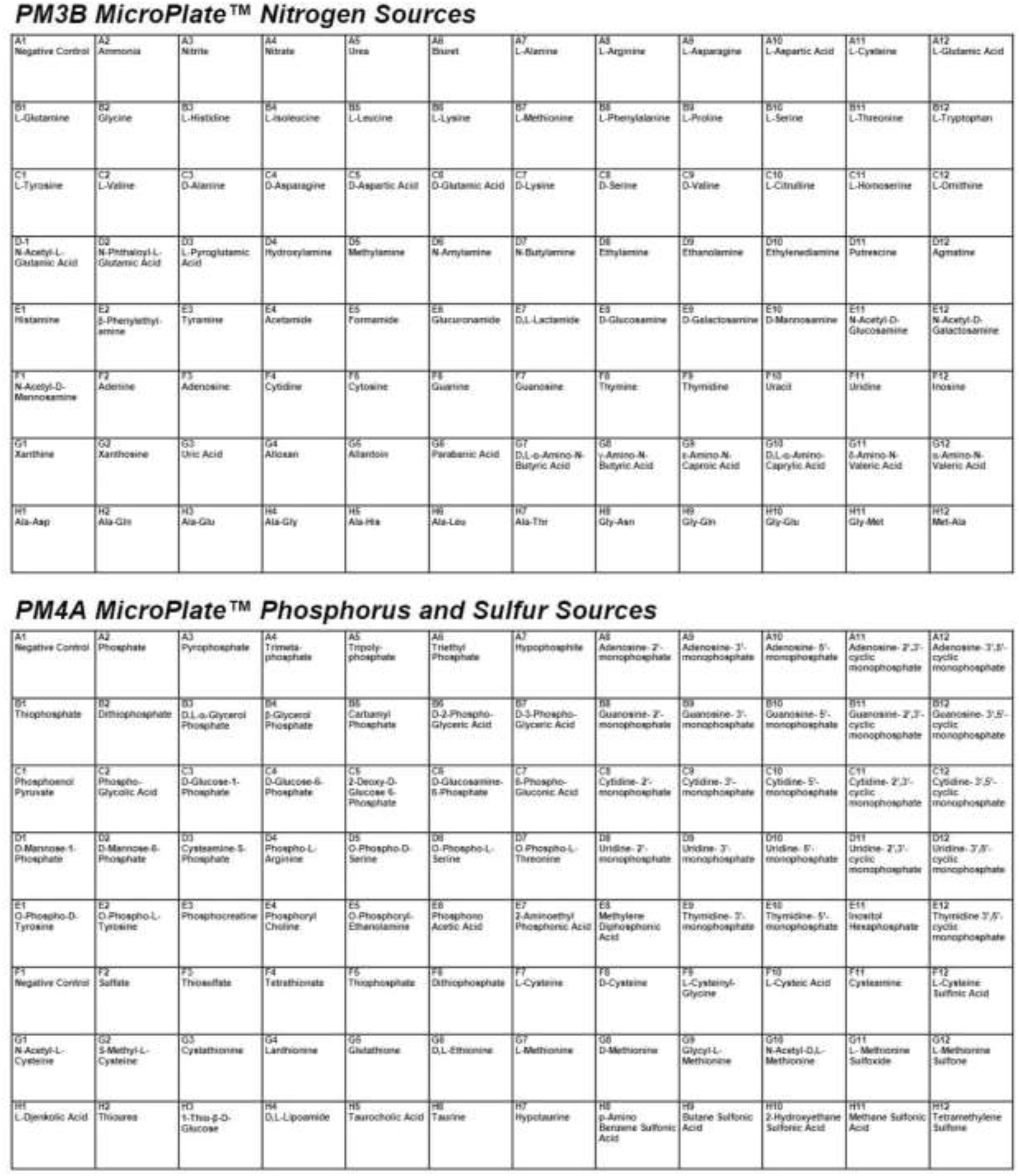
Composition of the compound arrays PM1, PM2A, PM3B and PM4A used for ligand screening.

**Fig. S2).**
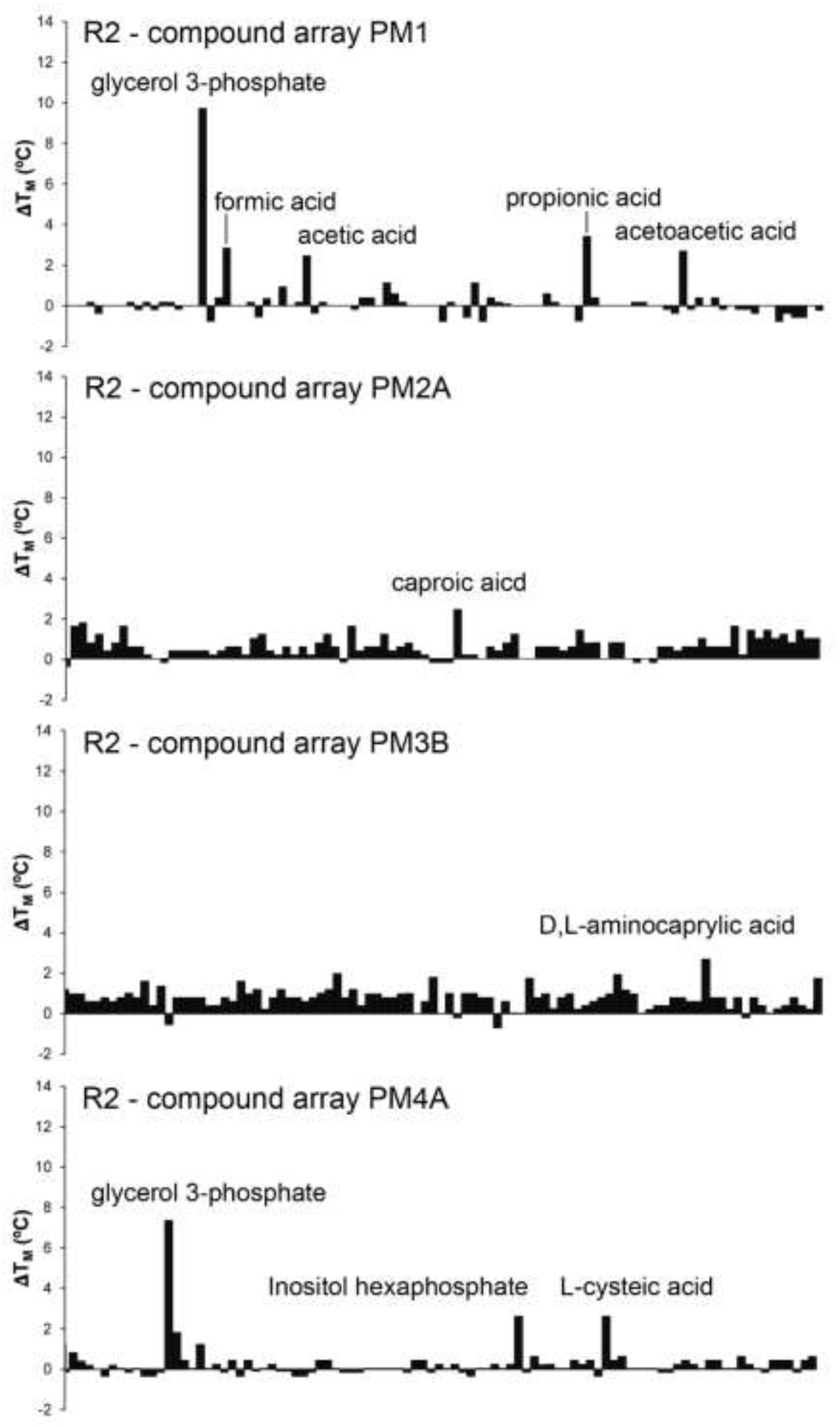
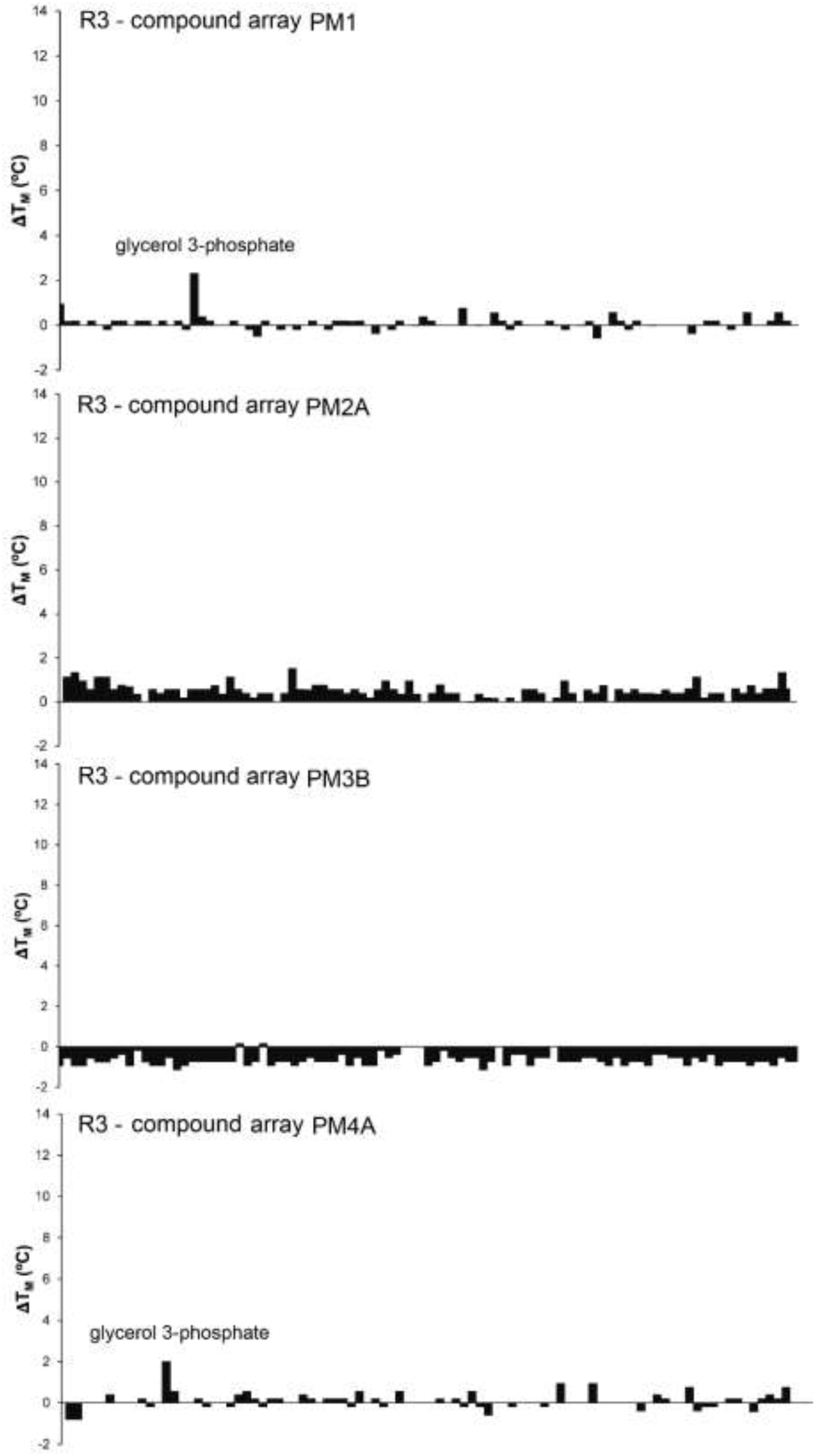
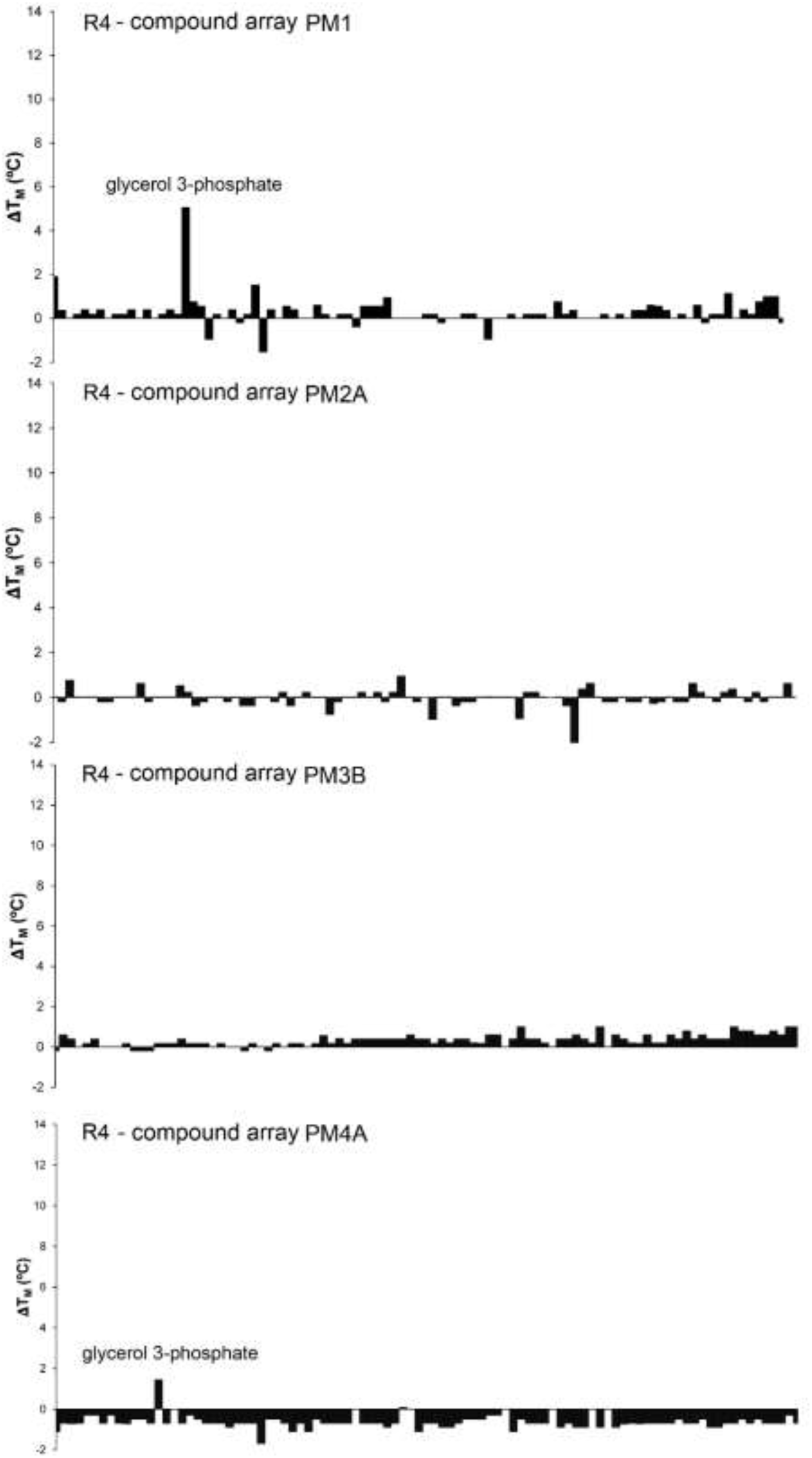
Changes in the midpoint of the protein unfolding transition (Tm) derived from thermal shift assays of domains R2, R3 and R4 with compound arrays PM1, PM2A, PM3B and PM4A (see Fig. S1 for composition) Microcalorimetric titrations of R2 with maximal possible concentrations of formic acid, acetic acid, propionic acid, acetoacetic acid, caproic acid, D,L-aminocaprylic acid and L-cysteic acid did not reveal binding.

**Table S1).**
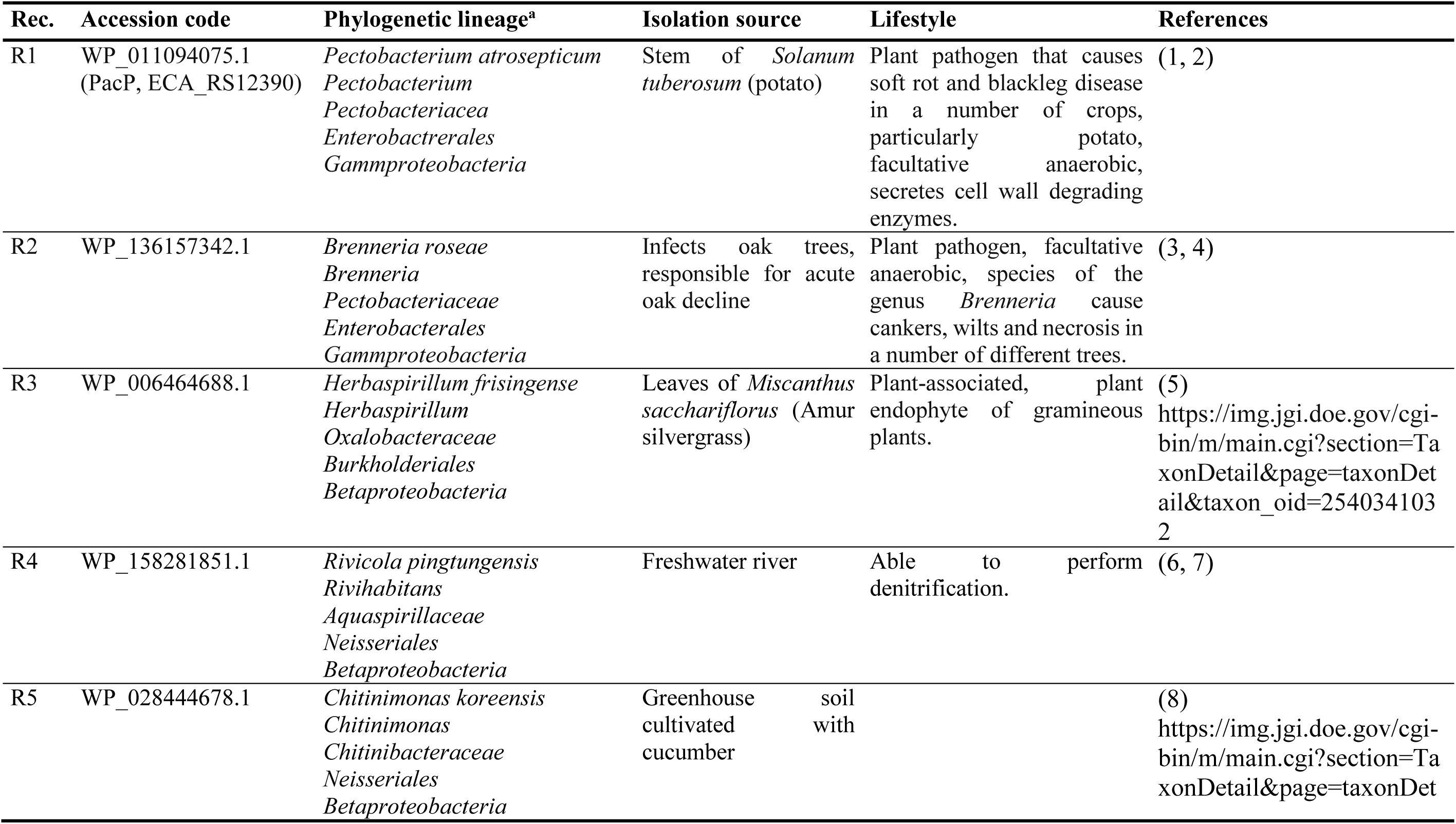
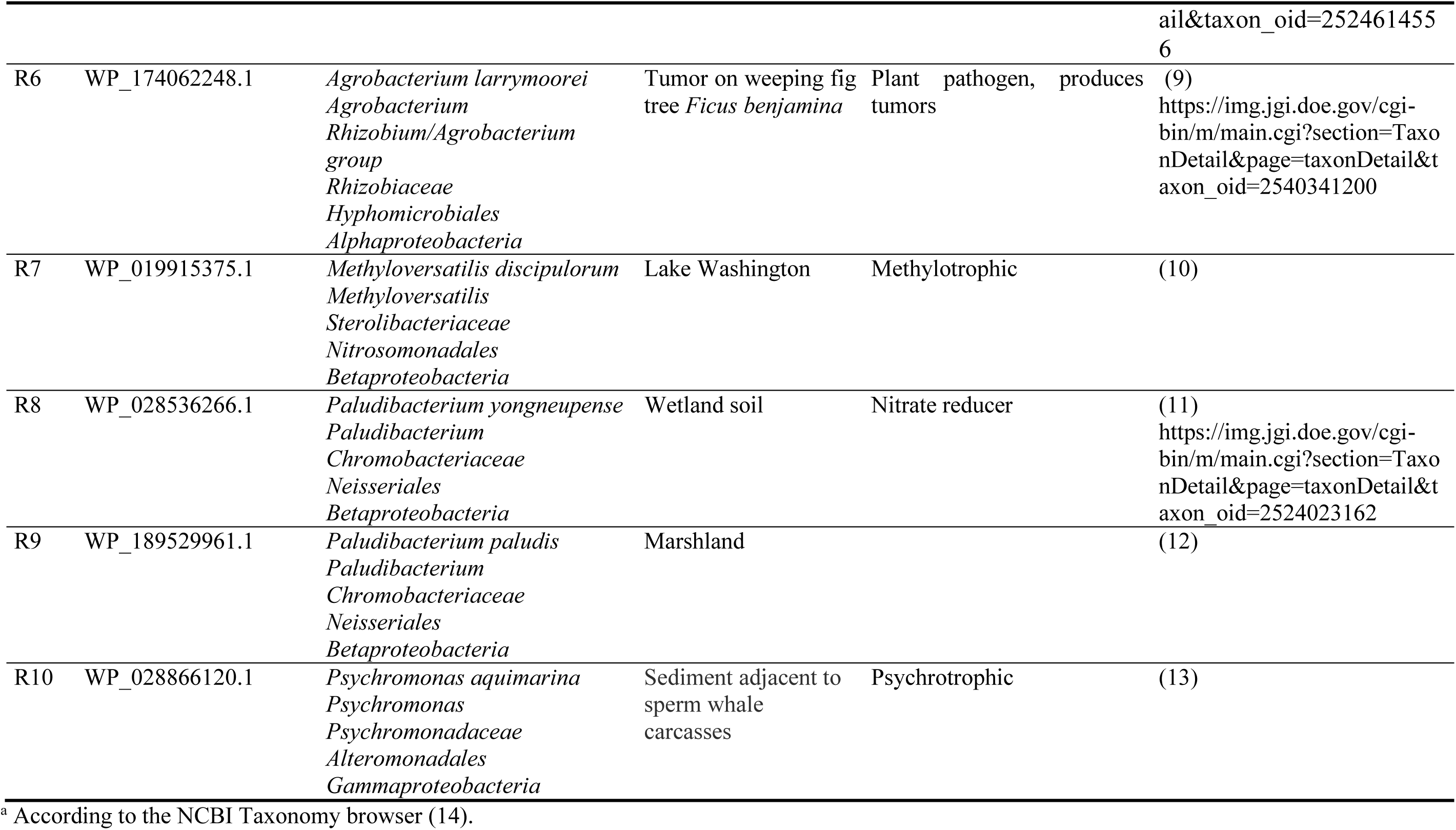
Information on the phylogenetics, lifestyle and isolation sources of strains containing chemoreceptors selected for study.

**Table S2).**
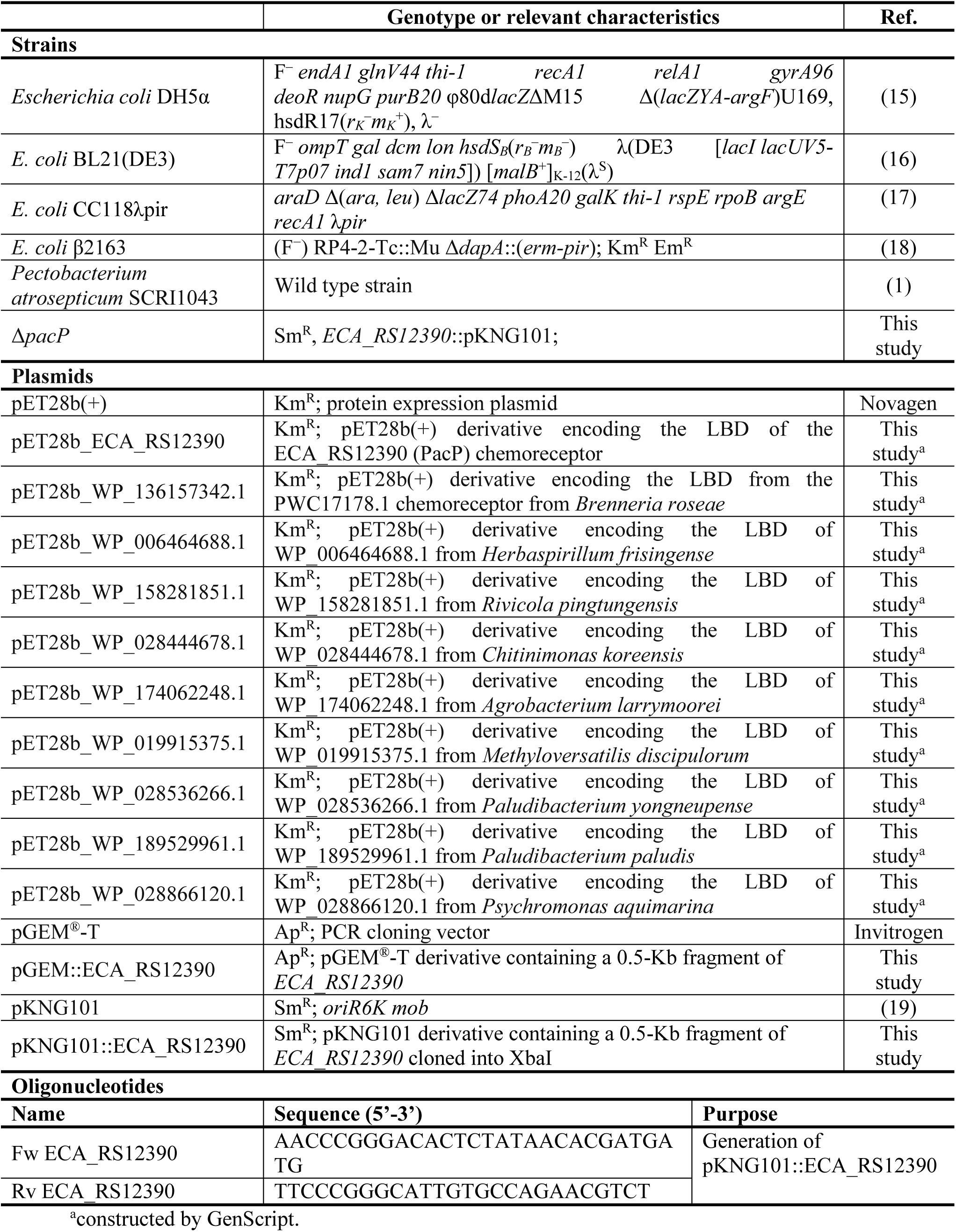
Strains, plasmids and oligonucleotides used in this study.

**Table S3).**
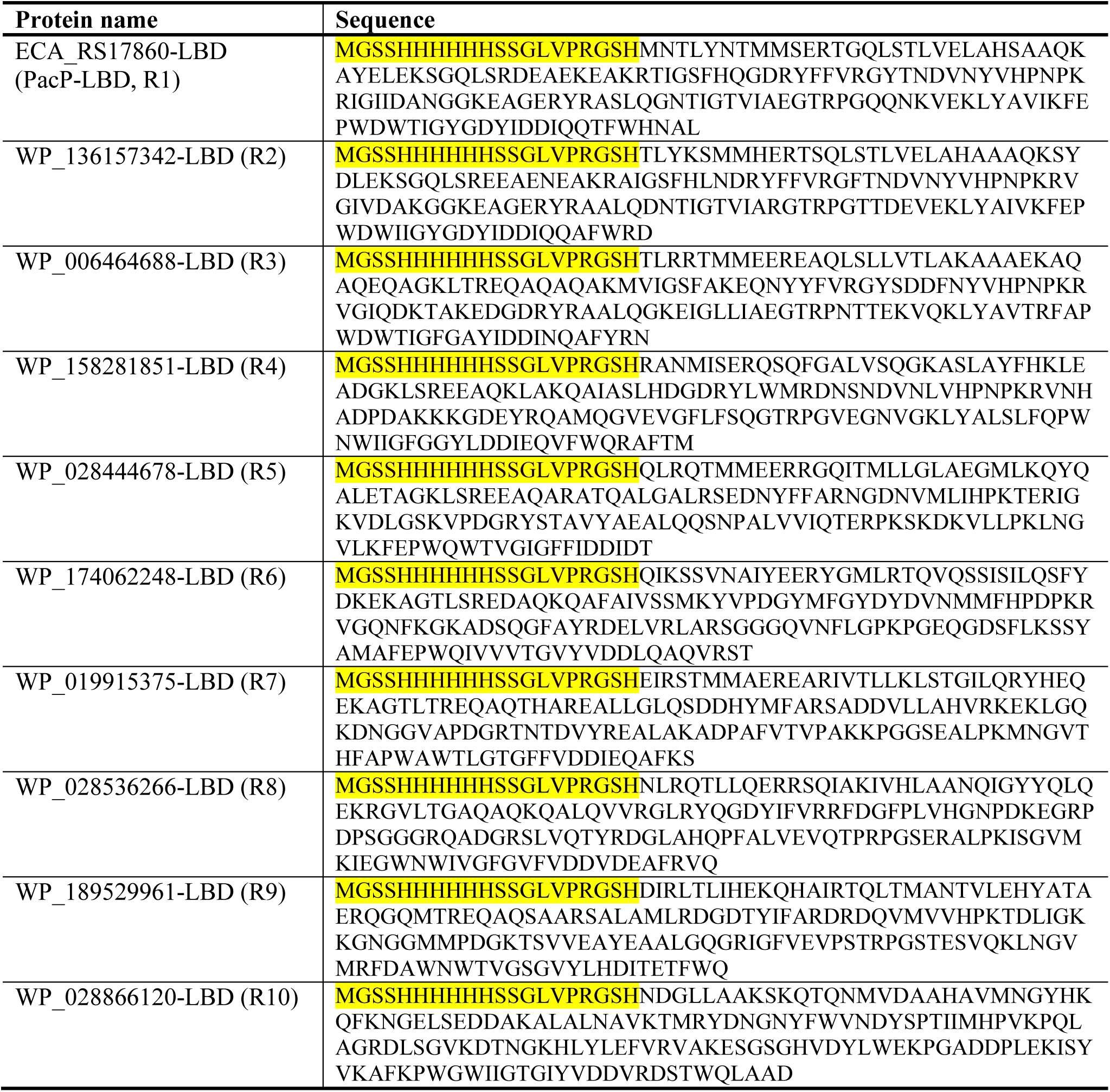
Sequences of proteins analyzed in this study. The C-terminal extension containing the his-tag is shaded in yellow.

**Table S4).**
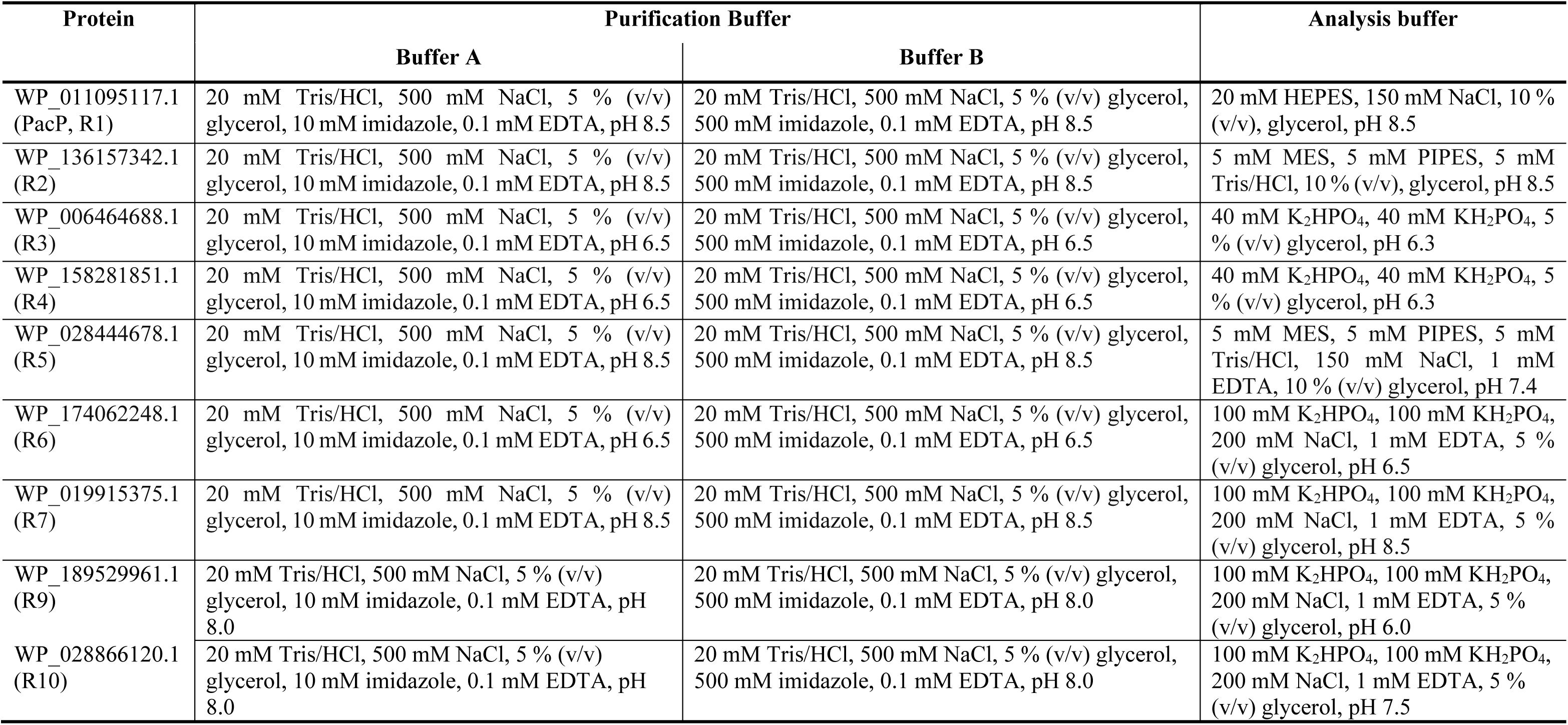
Buffers used for the purification and analysis of proteins. No expression in *E. coli* was observed for R8.

